# Unraveling the Molecular Complexity of Bicuspid Aortopathy: Lessons from Comparative Proteomics

**DOI:** 10.1101/2023.12.05.570304

**Authors:** Bárbara Pozo-Vilumbrales, Laura Martín-Chaves, Miguel A. López-Unzu, María Teresa Soto-Navarrete, Javier Pavón-Morón, Jorge Rodriguez-Capitán, Borja Fernández Corujo

## Abstract

**Background:** Molecular markers and pathways involved in the etiology and pathophysiology of bicuspid aortopathy are poorly understood. The aim here is to delve into the molecular and cellular mechanisms of the disease and identify potential predictive molecular markers using a well-established isogenic hamster model (T-strain) of bicuspid aortic valve (BAV) and thoracic aortic dilatation (TAD).

**Methods:** Comparative quantitative proteomics combined with western blot and morpho-molecular analyses in the ascending aorta of tricuspid aortic valve (TAV) and BAV animals from the T-strain, and TAV animals from a control strain. This strategy allows discriminating between genetic and hemodynamic factors in genetically homogeneous populations.

**Results:** The major molecular alteration in the aorta of genetically homogeneous BAV individuals is PI3K/AKT overactivation caused by changes in the EGF, ANGII and TGF-β pathways. PI3K/AKT affects downstream eNOS, MAP2K1/2, NF-κB, mTOR and WNT pathways. Most of these alterations are seen in independent patient studies with different clinical presentations, but not in TAV hamsters from T-strain that mainly exhibit WNT pathway downregulation.

**Conclusions:** We identify a combination of defective interconnected molecular pathways, directly linked to the central PI3K/AKT pathway, common to both BAV-associated TAD patients and hamsters. The defects indicate smooth muscle cell shift towards the synthetic phenotype induced by endothelial-to-mesenchymal transition, oxidative stress and inflammation. WNT signaling represent one genetic factor that may cause structural aortic abnormalities and aneurysm predisposition, whereas hemodynamics is the main trigger of molecular alterations, probably determining aortopathy progression. We identify twenty-seven novel potential biomarkers with a high predictive value.

## Introduction

Bicuspid aortic valve (BAV) is the most common congenital cardiac malformation, with an incidence of 1-2% in the general population^1^. BAV inheritance follows a complex pattern with low penetrance^2^. Thoracic aortic dilatation (TAD) occurs in up to 50% of BAV patients, who are at risk of aneurysm and dissection of the artery, which depicts the main mortality causes on BAV subjects^3^. Currently, preventive surgery stands as the only effective treatment for BAV aortopathy, with aortic diameter being the main reference criterion used to determine aortic dilatation and monitor the potential progression of the pathology^1^. Therefore, the search for circulating biomarkers in BAV aortopathy is an emerging area of research^4^.

The histopathological substratum of BAV aortopathy is cystic medial degeneration, which results from homeostatic alterations associated with dysregulation of TGF-β signaling pathway, abnormal stress mechanosensing and smooth muscle cell (SMC) phenotype switching^2,5^. Although multiple molecular and cellular pathways have been shown to be involved in disease etiology and progression, a unified mechanistic model is still missing.

The association between BAV and TAD is explained by two complementary hypotheses^6–8^. The hemodynamic hypothesis postulates that the BAV anatomy causes a turbulent blood jet stream that increases the stress on the aortic media, leading to deterioration of its histological structure. The mainstay of the second hypothesis is genetics. The defect leading to BAV during embryonic development may also entails a congenital structural abnormality in the ascending aorta, predisposing to aortic wall degeneration. Currently, most specialists favor the coexistence of hemodynamic and genetic factors in the disease etiology, although the exact influence of each factor remains unknown.

Our group has developed the only spontaneous animal model of BAV disease, which consists of an inbred (genetically uniform) strain (T-strain) of Syrian hamsters (*Mesocricetus auratus*) with a high incidence of isolated (non-syndromic) BAV^9–13^. Anatomy and inheritance of BAV in the hamster model are similar to those in patients^6,11,14–16^. The penetrance of the trait is incomplete, allowing the presence of both BAVs and tricuspid aortic valves (TAVs) in genetically uniform individuals^9,11,14–16^. Indeed, incidence of BAV in the T strain is the same in offspring of TAV animals as in offspring of BAV animals^9,11^, demonstrating that the coexistence of TAVs and BAVs results from incomplete penetrance of the trait and not from genetic variability. BAV in hamsters is caused by alterations in the septation of the embryonic cardiac outflow tract, due to defective behavior of the cardiac neural crest (CNC)^12^. Although T-strain animals do not exhibit conspicuous aneurysm of the aorta, both TAV and BAV individuals show TAD and express the histopathological substratum of the aortopathy irrespective of the aortic valve morphology, constituting experimental evidence supporting the genetic hypothesis on the etiology of BAV aortopathy^17^. This strain is well known and recognized as a reliable model for studying BAV disease including predisposition to aortopathy^1,4,8,12-^

In the present study, we have performed a comparative quantitative proteomic analysis of the structurally abnormal ascending aorta of hamsters with BAV (TBAV) and TAV (TTAV) from the T-strain, and TAV hamsters from an outbred strain (H strain) with no incidence of BAV (HTAV) and a normal aorta^17^. We have identified a set of interrelated altered molecular pathways common to BAV-associated TAD in hamsters that can be extrapolated to patients. In addition, we identified novel proteins that hold promise as predictive or diagnostic biomarkers. Our findings provide a unified perspective of the genetic and hemodynamic hypotheses of BAV aortopathy, uncovering molecular and cellular pathways distinctly affected by genetic and hemodynamic factors.

## Material and Methods

### Animals

The T-strain shows ∼40% incidence of BAV type A (2-sinus BAV anterior-posterior according to Michelena et al.^1^) and TAD^17^. The H strain (RjHan:AURA; Janvier, France) is a commercial outbred strain with 0% of BAV and TAD, previously used as a control group^17,18^.The handling protocol was approved by the Ethical Committee for Animal Experimentation of the University of Malaga (CEUMA; ethical authorization number: 25/08/2021/118).

A total of 44 adult animals (22 males, 22 females, ∼300 days old) were grouped according to the strain (T or H) and the aortic valve morphology (BAV or TAV): TBAV (n=19); TTAV (n=6); HTAV (n=19). The animals were euthanized using CO_2_ inhalation. The aortic valve morphology was evaluated, and the ascending aorta was excised and processed for protein extraction or histological procedures.

### Protein extraction, identification, and quantification (quantitative proteomics)

Each ascending aorta was cryopreserved in liquid nitrogen and stored at −80°C until use. A total of 18 specimens were included (TBAV: n=6; TTAV: n=6; HTAV: n=6). For each sample, two aortas were pooled and mechanically homogenized, followed by addition of 100 µl RIPA (Sigma-Aldrich) containing 0.5% protease inhibitor cocktail (Sigma-Aldrich). Finally, the samples were centrifuged at 13,000 rpm for 10 min at 4°C. Only the supernatant was retained. Protein concentrations were measured with a Qubit 4 Spectrofluorometer (Thermo Fisher). Three biological replicates, each with 50 µg of protein, were used for the proteomic analysis.

The identification and quantification of peptides were carried out following the previously described method^19^ (detailed in the supplementary material). In brief we used an Easy nLC 1200 UHPLC system coupled to a linear quadrupole E-trap hybrid mass spectrometer-Orbitrap Q-Exactive HF-X (Thermo Fisher). Data were acquired using Tune 2.9 and Xcalibur 4.1.31.9 i. Survey mass range was m/z 375-1.600 at a resolution of 120.000. MS/MS2 spectra searches were performed against the NCBI protein database of *M. auratus*. Protein quantification was implemented by label-free.

### Western blot and Immunofluorescence

To validate and extend key proteomic findings, some proteins were analyzed by western blot (WB) and immunofluorescence (IF) as previously described^17,19^, and detailed in the supplementary material.

Six specimens (TBAV: n=3; HTAV: n=3) were used for WB, and 10µg of proteins were analyzed by SDS/PAGE.

Regarding IF, 10 animals were included (TBAV: n=5; HTAV: n=5). Ascending aortas were fixed by overnight immersion in 4% paraformaldehyde in PBS, dehydrated in graded ethanol, and embedded in paraffin (Histosec; Merck). Eight µm serial transverse sections were used to localize and quantify α-Actin expression. Nuclei were stained with 4′,6-diamidino-2-phenylindole (DAPI) and images were obtained with a Leica SP8 confocal microscope.

The primary antibodies used were polyclonal anti-pAKT (Cell Signaling), polyclonal anti-pSMAD2 (Thermo Fisher) and monoclonal anti-α-Actin (Sigma Aldrich).

### DAF-2DA

For nitric oxide (NO) detection, ascending aorta of 10 animals (TBAV: n=5; HTAV: n=5) were incubated in 4,5-diaminofluorescein diacetate (DAF-2DA; Sigma-Aldrich) as previously described^20^. Briefly, samples were incubated for 12h at 37°C in 10µM DAF-2DA diluted in Tyrode’s buffer or in Tyrode’s buffer only as a negative control. Then, tissues were fixed in paraformaldehyde 4% in PBS overnight and embedded in Tissue-Tek (Sakura Finetek) for cryosectioning. Transversal sections were cut at eight μm in a Leica CM3050S cryostat, and nuclei were stained with DAPI. Images were obtained with a Leica SP8 confocal microscope. Due to the autofluorescence of aortic lamellae, the baseline fluorescence was stablished with the negative control. The quantification methos is detailed in supplementary material.

## Results

We identified 2265 proteins in TBAV and HTAV aortas. Of these, 2115 (93.4%) were present in both groups (Fig.1A). From the remaining 150 proteins, 19 were exclusively detected in every replicate of HTAV and eight of TBAV animals (Supplemental Table 1). A total of 111 proteins were differentially expressed in TBAV compared to HTAV individuals, 40 proteins were upregulated and 71 downregulated in the TBAV group (Fig.1B,C; Supplemental Table 2). This set of proteins showed a protein-protein interaction (PPI) enrichment p-value=4.93 E-12 in the STRING database, meaning that they interact more than expected in a set of random proteins (Fig.2A).

**Figure 1.**
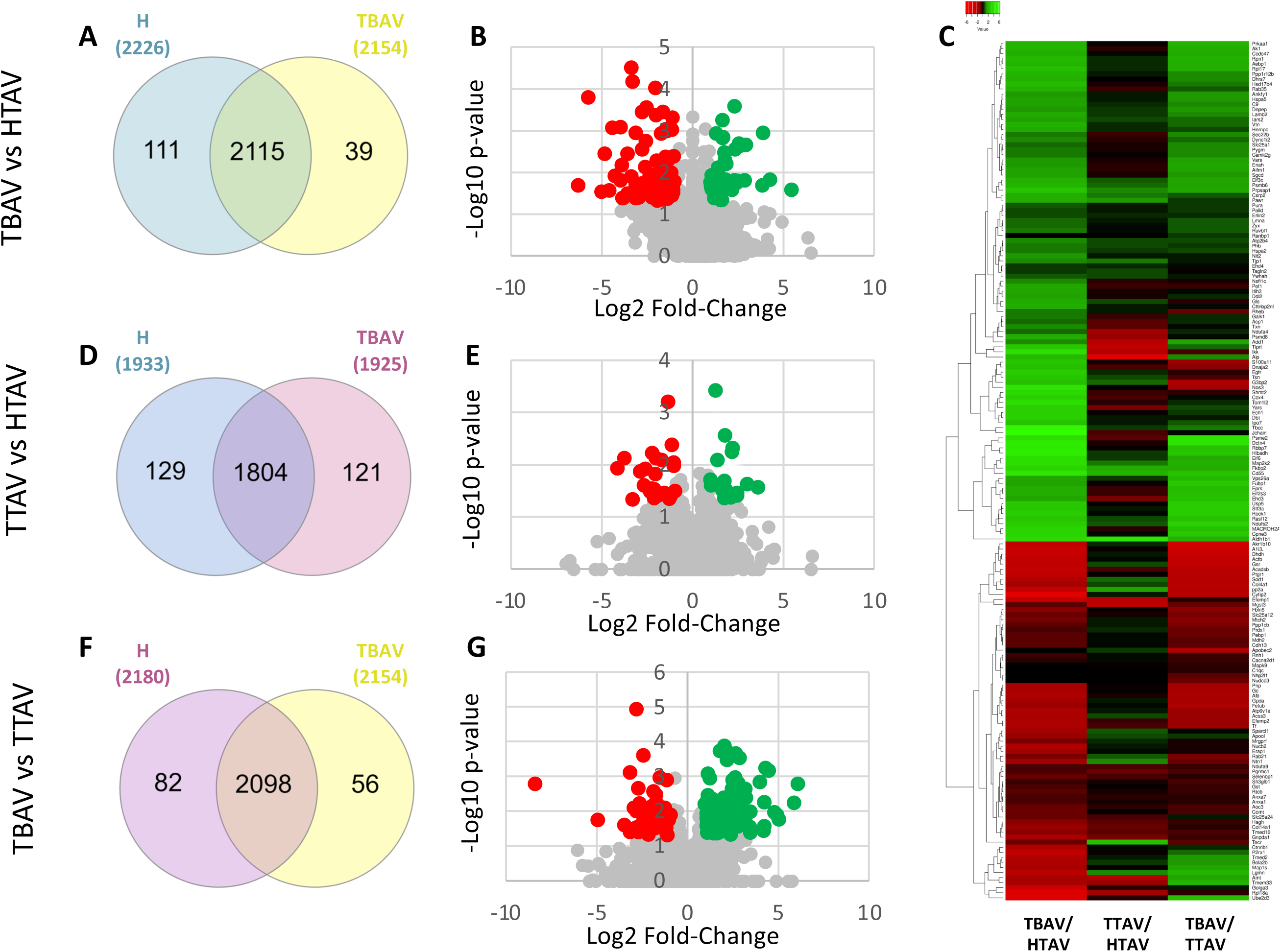
Proteomic analysis of the ascending aorta between experimental groups. (A,B) TBAV *vs.* HTAV. (D,E) TTAV *vs.* HTAV. (F,G) TBAV *vs.* TTAV. (A,D,F) Venn diagram. (B,E,G) Volcano plots. (C) Heatmap. For B,C,E,G significantly downregulated and upregulated proteins are labeled in red and green respectively.

**Figure 2.**
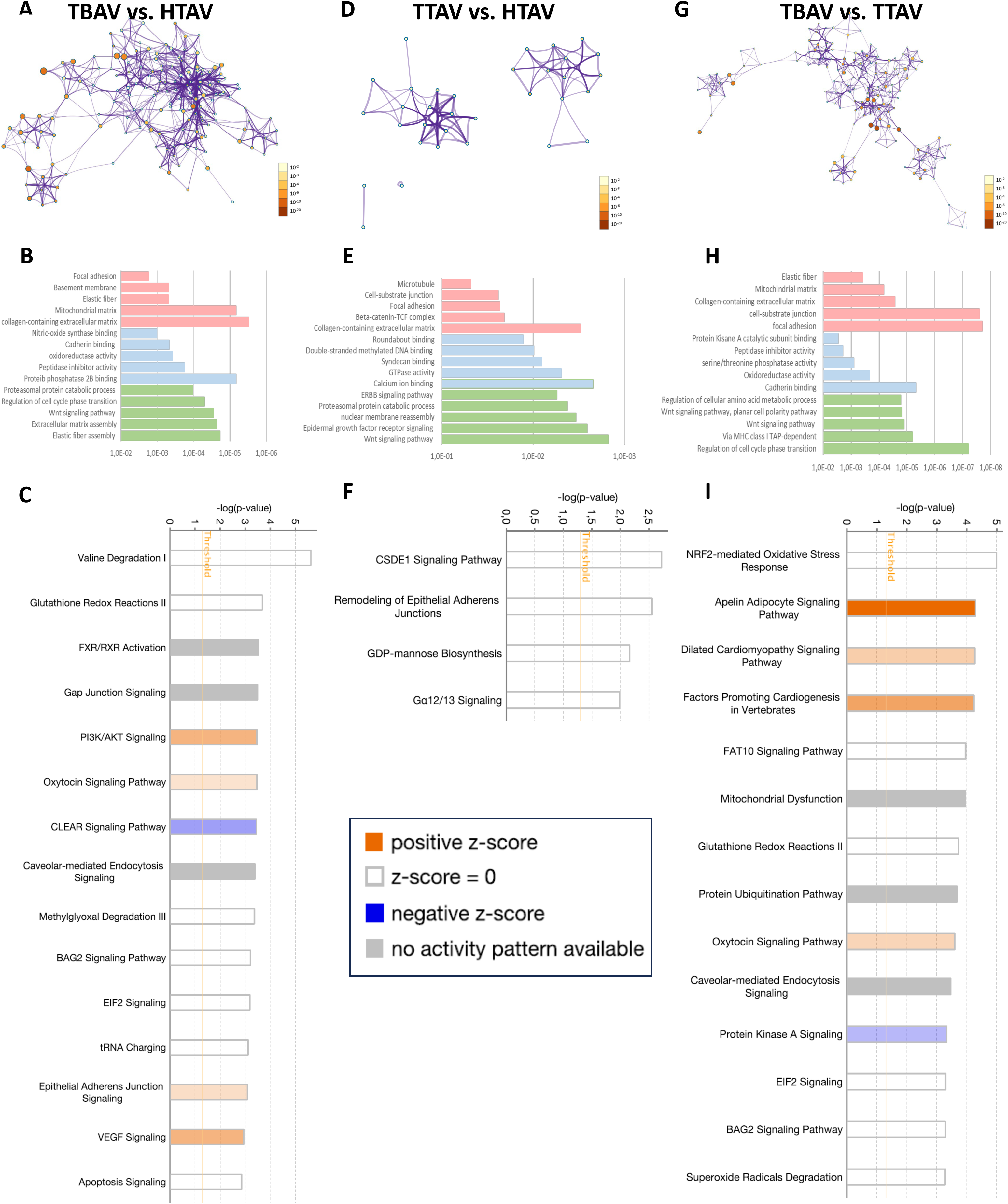
Enrichment analysis of differentially expressed proteins. (A-C) TBAV *vs.* HTAV; (D-F) TTAV *vs.* HTAV and (G-I) TBAV *vs.* TTAV. (A,D,G) Protein-Protein interaction (PPI) network performed with Metascape. Nodes represent proteins, and edges protein-protein associations. (B,E,H) Overview of enrichment analysis based on protein alterations among experimental groups. Green: biological processes; blue: molecular functions; pink: cellular components. (C,F,I) Top 15 altered canonical pathways identified by IPA.

The functional annotation of proteins based on GO terms is shown in figure 2B. According to the enrichment factor, the most representative affected biological processes (p-values<1.05E-4) were elastic fiber assembly, extracellular matrix assembly, WNT signaling pathway, regulation of cell cycle phase transition and proteasomal protein catabolic process. The most significantly affected molecular functions (p-values<1.01E-3) were protein phosphatase 2B binding, peptidase inhibitor activity, oxidoreductase activity, cadherin binding and nitric-oxide synthase binding. The most representative cellular components (p-values<1.72E-3) were collagen-containing extracellular matrix, mitochondrial matrix, elastic fiber, basement membrane and focal adhesion.

The analysis of differentially expressed proteins with the ingenuity pathway analysis (IPA) software showed 51 enriched canonical pathways according to the −log_10_ (p-value) >|1| threshold (Supplemental Table 3). Of these, 14 pathways showed a Z-Score >|1|, where the upregulation of PI3K/AKT pathway exhibited the highest statistical significance and Z-Score (p-value=3.38 E-4; Z-Score=1.00) (Fig.2C). The IPA causal network analysis identified 15 regulatory molecules showing causal relationships with our data (Supplemental Table 4). ATR-1 had the highest activation prediction (p-value=7.07 E-05; Z-Score=2.496).

To validate the quantitative proteomic results, the expression levels of several key proteins were assessed in HTAV and TBAV animals. WB showed that pSMAD2, pAKT and eNOS were upregulated, whereas α-Actin was downregulated in TBAV compared to HTAV animals (Fig.3A). IF demonstrated a reduction of α-Actin^+^ SMCs in the aortic media of TBAV compared to HTAV animals (Fig.3B-D). In addition, the abundance of NO, assessed by DAF-2DA, exhibited a drastic decrease in the aorta of TBAV animals (Fig.3E-G).

**Figure 3.**
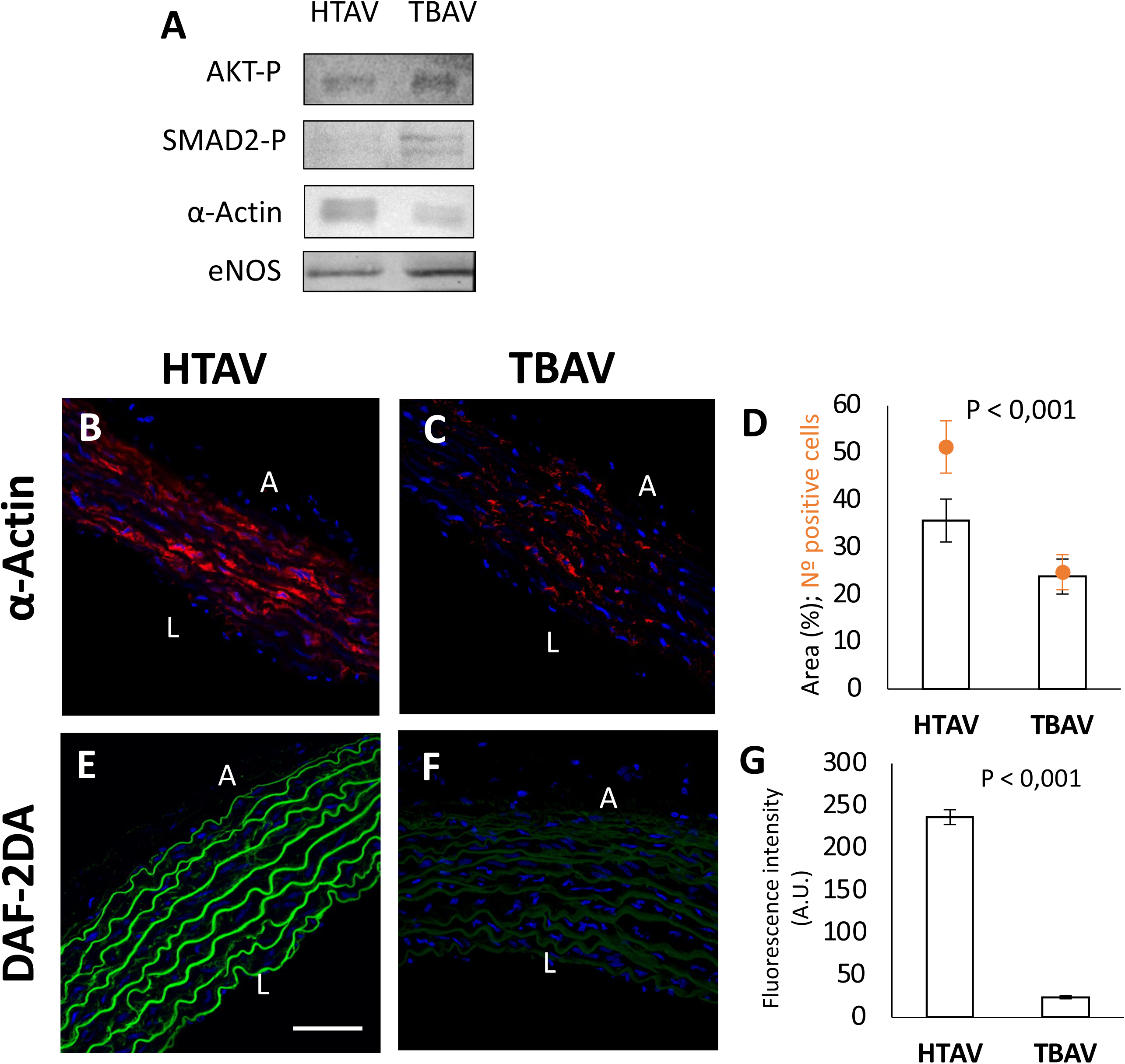
Validation of proteomics results. (A) WB of pAKT, pSMAD2, α-Actin and eNOS. (B-D) Immunofluorescence of α-Actin (red) in HTAV (B) and TBAV (C) aortas, quantification of the area occupied by positive cells (D, bars) and of the number of positive cells (D, orange). (E-G) DAF-2DA fluorescence (green) in HTAV (E) and TBAV (F) aortas, and quantification of intensity (G). Nuclei were stained with DAPI (blue). A: adventitia, L: lumen. Scale bar: 50 μm.

Proteomic comparisons between TTAV and HTAV revealed 2054 proteins in the ascending aorta, of which 1804 (87.8%) were present in both groups (Fig.1D). Two unique proteins were detected in each replicate of the HTAV group, and three unique proteins in the TTAV group (Supplemental Table 5). The volcano plot (Fig.1E) and the heatmap (Fig.1C) showed 27 differentially expressed proteins (11 upregulated and 16 downregulated in TTAV animals; Supplemental Table 6), with a PPI enrichment p-value=0.358, meaning that they do not form a specific set of interacting proteins (Fig.2D). Based on the GO terms enrichment analysis (Fig.2E), the most representative affected biological processes (p-values<0.0054) were WNT signaling pathway, EGFR signaling pathway, nuclear membrane reassembly, proteasomal protein catabolic process and ERBB signaling pathway. The most significantly altered molecular functions (p-values<0.0127) were calcium ion binding, GTPase activity, syndecan binding, double-stranded methylated DNA binding and roundabout binding. The most representative affected cellular components (p-values<0.048) were collagen-containing extracellular matrix, beta-catenin-TCF complex, focal adhesion, cell-substrate junction, and microtubule. The figure 2F shows the four only pathways that the IPA software found to be significantly altered (p<0.001; Supplemental Table 7), where the Cold Shock Domain-containing E (CSDE1) pathway showed the highest significance, although none of them had an associated z-score.

Finally, comparison between TBAV and TTAV animals revealed 2236 proteins. Of these, 2098 proteins (93.8%) were present in both groups (Fig.1F). Three unique proteins were identified in all replicates of the TBAV group and 14 in the TTAV group (Supplemental Table 8). The volcano plot (Fig.1G) and the heatmap (Fig.1C) revealed 110 altered proteins (77 upregulated and 31 downregulated in TBAV animals) with a PPI enrichment p-value=5.59 E-4 (Fig.2G; Supplemental Table 9), indicating that these proteins, as a group, have a significant biological association.

The enrichment analysis of GO terms (Fig.2H) indicated that the most representative altered biological processes (p-values<01.54E-5) were regulation of cell cycle phase transition, via MHC class I TAP-dependent, WNT signaling pathway, WNT signaling pathway planar cell polarity pathway and regulation of cellular amino acid metabolic process. The most significantly altered molecular functions (p-values<1.99E-3) were cadherin binding, oxidoreductase activity, calcium-dependent protein serine/threonine phosphatase activity, peptidase inhibitor activity and protein kinase A catalytic subunit binding. The most representative altered cellular components (p-values<3.83E-4) were focal adhesion, cell-substrate junction, collagen-containing extracellular matrix, mitochondrial matrix and elastic fiber.

Based on the IPA analysis, the proteins with altered expression were associated with 50 enriched canonical pathways (Supplemental Table 10). Topmost significantly altered pathways, with p-values<5.12 E-5 and Z-Scores >|1|, were Apelin Adipocyte Signaling pathway, Dilated Cardiomyopathy Signaling Pathway and Factors Promoting Cardiogenesis in vertebrates (Fig.2I).

## Discussion

The T strain of Syrian hamsters constitutes a consolidated spontaneous animal model of non-syndromic congenital BAV^1,3,4,8–10,12,13,17,18^. In the present study, we found 138 differentially expressed proteins in the aorta of TBAV animals compared to animals from a control strain. A significant proportion of these proteins are involved in human BAV aortopathy or have molecular and/or cellular functions that align with the proposed mechanisms of disease (Fig. 2). Thus, our results confirm the utility of the T-strain as a model of TAD and bicuspid aortopathy predisposition.

Compared to other proteomic studies including aortic tissue from patients^21,22^, the number of proteins identified, their concordance among replicates, and their homology among experimental groups demonstrate a high degree of consistency. This reliability is explained by the genotypic (inbred, isogenic) and phenotypic (non-stenotic, antero-posterior BAV) homogeneity of the animal model^9,10,12,17,18^, meaning a strong advantage for comparative analysis with respect to studies in patients. The latter usually include populations with different clinical presentations associated with different disease etiologies, which limits the stringency of the results, whereas our hamster population includes animals with BAV type A exclusively, with a specific etiology. This is relevant for the interpretation of the present results, and their extrapolation to patients.

During the last decade, several key molecular pathways have been found to contribute to aneurysm formation, notably TGF-β, WNT, NOTCH, PI3K/AKT and ANGII, supported by numerous independent evidence^23–27^. However, although both experimental and human studies support the involvement of these pathways, it is recognized that the evidence base remains fragmentary. The key missing question, as experts have already pointed out^24^, is whether these pathways are functionally interconnected in the ascending aorta, such that the combination of their alterations causes cellular dysfunction, ultimately leading to tissue deterioration. Alternatively, different molecular pathway alterations may characterize an etiologically distinct patient population. Given that the hamster model represents a specific etiology, we expected to obtain a restricted set of altered proteins associated with a specific pathway. Remarkably, we found that four of these five major pathways were interconnected and concordantly altered in our TBAV individuals (Fig.4). Thus, we show for the first time that 1) this set of defective pathways, independently implicated in human BAV aortopathy cross-talks in the same ascending aorta; 2) given that this set appears both in heterogeneous patient cohorts and in a homogeneous animal model, we can deduce that there exists a specific assemblage of defective molecular pathways shared by most patients and hamsters with BAV, regardless of their etiology. This is a key finding of the present study that frames the interpretation of the specific results.

**Figure 4.**
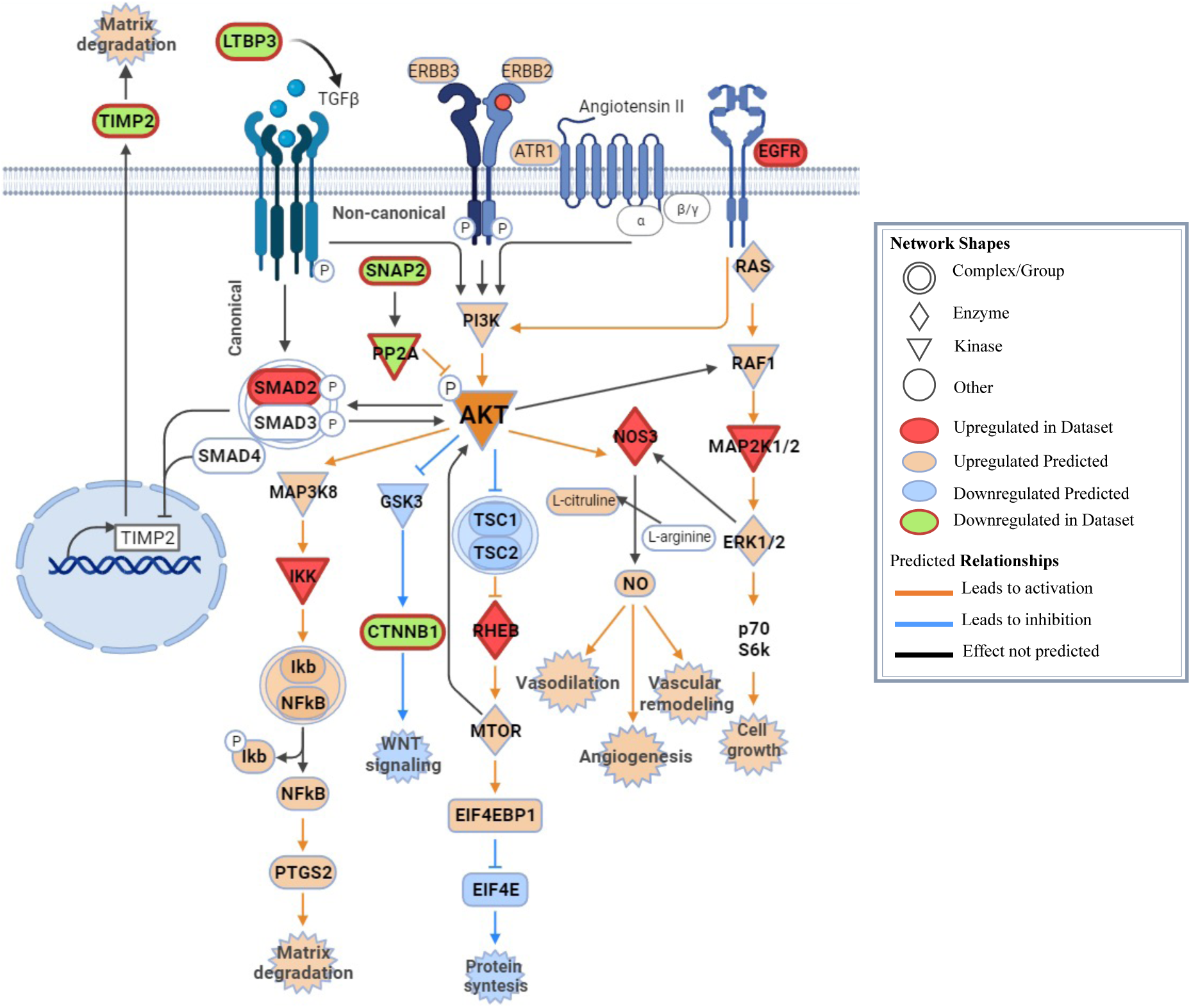
Crosstalk regulation between EGFR, ATR-1, TGF-β and AKT signaling. The diagram is built from the TBAV *vs*. HTAV IPA analysis, after adding the unique proteins in TBAV. PI3K/AKT signaling pathway was the most significantly affected in TBAV animals. See the labelling in the Prediction Legend.

Our proteomic analysis indicates that PI3K/AKT signaling is a central pathway within this assemblage. WB results confirm the upregulation of pAKT in TBAV animals. In our dataset, this pathway exhibits the most significant alterations, with five upregulated (SMAD2, IKK, RHEB, MAP2K1/2, and eNOS) and four downregulated (LTBP3, SMAP2, PP2A, and CTNNB1) proteins in TBAV hamsters (Fig.4). Notably, most of these proteins are independently altered in different patient populations^8,23,24,26,28,29^. In SMCs of some BAV patients, the overactivation of PI3K/AKT pathway results from an increase of pAKT, regardless of the aortic diameter, suggesting a congenital predisposition of these patients^30^. This hypothesis aligns with our animal model, in which molecular and structural alterations of the ascending aorta are associated with BAV morphology in the absence of aneurysm, constituting a model of predisposition^17^.

Both the GO terms and IPA analyses of the proteomic outcome revealed significant alterations of cellular processes and canonical pathways directly associated with PI3K/AKT signaling (Fig.2B,C; Supplemental Table 3), which indicate dysfunctions both upstream and downstream of this pathway in the aorta (Fig.4).

The detected altered PI3K/AKT upstream pathways include the TGF-β canonical and non-canonical pathways, EGF and ANGII pathways. Involvement of the former is supported by detection of increased activated SMAD2 (Fig.3A). In both patients and T-strain, alterations of the TGF-β pathway constitute the most persistent mechanism involved in BAV-associated TAD^7,17,23,31^. Dysregulation of this pathway may cause TIMP downregulation in hamster (Fig.4) and in patients^32^ promoting matrix degradation. EGF is an activator of both the PI3K/AKT and MAPK pathways that regulate the eNOS pathway^33^. We found upregulation of EGFR in our analysis (Fig.4). This is the first time that the EGFR pathway is proposed to play a role in TAD, what seems plausible given the reported activation of EGFR upon mechanical stress caused by turbulent flow^34^. IPA predicted the upregulation of ATR-1, one of the ANGII receptors. This protein could mediate an increase in TGF-β levels in patients with BAV-associated TAD, leading to SMC proliferative phenotype and ECM remodeling^25^. Therefore, overactivation of EGFR, TGFBR and ATR-1 signaling would have an additive effect on the regulation of the PI3K/AKT activation pathway, positioning it as a central pathway for BAV aortopathy (Fig.4). Additionally, we detected a significant downregulation of SMAP2 and PP2A, known inhibitors of AKT function^35,36^ (Fig. 4). PP2A is known to participate in the progression of abdominal aorta aneurysm^24^, and its pharmacological activation by SMAP2 has been proposed as a therapeutic strategy^37^.

Based on the IPA analysis, the PI3K/AKT upregulation in TBAV animals affects at least five pathways downstream: NF-κB, WNT/CTNNB1, mTOR, MAPK1/2, and eNOS (Fig.4). All these molecular pathways participate in SMC phenotype shift between contractile and synthetic phenotypes, a cellular process that occurs during human TAD. Normally, SMCs appear as not terminally differentiated, retaining a high degree of cellular plasticity, what allows changing their phenotype in response to severe environmental signals^38^. This contributes to the growth, remodeling, and repair of the vasculature, and also to the progression of cardiovascular pathologies^39^. During ascending aorta dilatation, SMC phenotype shift has been associated with an increase in oxidative stress and proteolytic enzyme production, what may contribute to degrade the ECM, and facilitate SMC detachment, migration, and apoptosis^40–43^. The pathways known to regulate SMC phenotype switching are GSk3b/CTNNB; RAS/RAF/MEK/ERK; NOTCH; PI3K/AKT and TGF-β^44^. Remarkably, we found dysregulation of four of these pathways in TBAV animals, attributed to the altered expression of nine proteins in those pathways (Fig.4). LTBP3, TGF-β, EGFR, SMAD2, REHB and MAPK1/2, involved in downregulation of contractile proteins^24^ were found upregulated, whereas GSK3b and β-catenin, involved in the differentiation of SMCs to the contractile phenotype^45^ were downregulated. These alterations are concordant with SMC shifting towards the synthetic phenotype, which was supported by the reduced number of α-Actin^+^ aortic SMCs in TBAV animals (Fig.3A-D).

TBAV animals showed a significant decrease in CTNNB1, which is consistent with the overactivation of the PI3K/AKT pathway (Fig.4). WNT/CTNNB1 serves as a master regulator of the epithelial-to-mesenchymal transition (EMT) process, which is responsible for the formation of the embryonic valve primordia^13^ and promotes the synthetic phenotype in adult aortic SMCs^29^. Additionally, WNT/CTNNB1 signaling regulates the migration of CNC cells into the embryonic cardiac outflow tract, to finally differentiate into valvular interstitial cells and aortic SMCs^13,46^. Alterations in both EMT and CNC migration have been proposed as causes of BAV and abnormal aortic development in patients, hamsters and several mutant mouse models^8,13,17,29^. In line with this result, we have also detected upregulation of RHEB in TBAV specimens, which regulates AKT translational responses through mTOR (Fig.4). Hyperactivation of mTOR signaling has been reported in BAV patients with aneurysm^47^. Intriguingly, this alteration contributes to a reduction in cell contractility and phenotype shift specifically in CNC-derived SMCs^8^. IKK, a central regulator of the NF-κB pathway (Fig.4)^48^, was significantly upregulated in TBAV animals, and the activation of this pathway has been described in TAD patients^49^. NF-κB regulates inflammatory responses, which favors SMC shift^50^. Our analysis revealed a significant eNOS upregulation, probably as a response to PI3K/AKT and MAP2K1/2 overactivation (Fig.4)^51^. However, we detected a significant decrease in the production of NO in the aortic media of TBAV animals (Fig.3E-G). This discrepancy can be explained by eNOS uncoupling, a process in which overexpression of the enzyme increases oxidative stress and decreases NO production, as detected with DAF-2DA (Fig.3E-G), ultimately inducing SMC phenotypic shift and apoptosis^51–55^. MAP2K1/2 overactivation may affect the Ras/Raf/MEK/ERK signaling pathway, which regulates the suppression of SMC specific mature markers and induces phenotype switching^44^. In addition, previous studies in Fbn1-null mice with accelerated aneurysm growth showed MAPK overactivation^56^. Our previous studies in the T-strain detected a decrease of Fbn1/Fbn2 expression ratio in the aorta of BAV animals^17^. This alteration might be involved in the increase of MAPK signaling, which would promote the non-canonical TGF-β pathway by promiscuous SMAD2 activation^56^. Interestingly, CNC development is critically dependent on the MAPK signaling pathway^57^.

Thus, the present proteomic analysis points not only to pathways associated with SMC phenotype shift, but also to CNC migration and differentiation. Four of the five main altered pathways detected in this study (WNT, mTOR, Ras/Raf/MEK/ERK, eNOS; Fig. 4) regulate CNC cell migration in the embryo and differentiation to SMC in the adult^43,44^. Taken together, all these findings support the notion that CNC-derived SMC exhibit abnormal shift to the synthetic phenotype, driven by WNT/CTNNB1-induced EMT and sustained by the mTOR pathway. NF-κB-induced inflammatory response and oxidative stress caused by eNOS uncoupling may favor the shift, together with alterations of TGF-β and MAPK1/2 pathways (Fig.4).

All the results presented above were obtained comparing BAV animals from the affected T-strain (TBAV) with control animals (HTAV). TBAV and HTAV animals differ in the aortic valve morphology and the genetic background, which contains still unidentified genetic variants responsible for TAD and BAV development in the T-stain. Therefore, aorta proteome differences between TBAV and HTAV animals are based on both, structural abnormalities caused by the genetic background and possible hemodynamic changes due to the bicuspid morphology of the aortic valve. TTAV and HTAV animals also differ in the genetic background, but it causes only TAD and not BAV in TTAV animals due to the reduced penetrance of the BAV trait^9,11,17,18^. Consequently, aorta proteome differences between TTAV and HTAV animals are based exclusively on structural abnormalities caused by the genetic background. Hemodynamic changes causing additional differential protein expression can be discarded, as tricuspid valve morphology is virtually identical in HTAV and TTAV animals^17,18^. When we compared TTAV and HTAV aortas, as few as 27 proteins showed a significant differential expression, of which ten were associated with the WNT/CTNNB1 pathway (Supplemental Table 6). Accordingly, the GO terms enrichment analysis indicated that WNT signaling is by far the most significantly altered pathway in this comparison. In addition, the IPA analysis revealed CSDE1, a direct regulator of WNT signaling^58^, as the most significantly altered pathway. These results indicate that protein changes in the aorta of TTAV animals are almost exclusively dominated by the WNT/CTNNB1 pathway. It must be deduced that alterations of the WNT/CTNNB1 pathway are exclusively associated with structural defects of the aorta, excluding the possible hemodynamic effect of the BAV morphology. As explained above, WNT signaling is a central regulator of two key embryonic processes involved in BAV formation, CNC cell migration and EMT, the latter also considered a strong stimulator of SMC phenotype shift in the adult aorta. The morphogenetic defect that gives rise to BAV type A in the hamster model is an alteration of CNC cell migration, inducing the excessive fusion of conotruncal ridges during valvulogenesis, probably mediated by EMT^12,13^. CNC cells differentiate into a proportion of the SMCs that build the adult aortic wall^25^. We hypothesize that an alteration in the WNT pathway could be responsible for the defective behavior of CNC cells in animals from T-strain, leading to BAV during embryonic development, and favoring the acquisition of the synthetic phenotype in CNC-derived SMCs of the adult ascending aorta. Therefore, WNT may constitute a central pathway supporting the genetic hypothesis on the etiology of BAV aortopathy, as alterations in this pathway can explain the etiological connection between embryonic BAV formation and adult aortic pathology.

To recapitulate, the TBAV *vs.* HTAV aortic proteome comparison results in a broad collection of altered pathways including WNT/CTNNB1, whereas TTAV *vs.* HTAV comparison results almost exclusively in alterations of the WNT/CTNNB1 pathway. Thus, it appears that, except for WNT, the diversity of altered pathways detected in TBAV animals is strictly associated with the BAV phenotype. It can be deduced that the valve morphology and not genetics is the trigger of most proteome alterations in TBAV animals. Hemodynamics associated with the valve morphology would be the most reasonable factor causing these changes. In fact, when we compared the proteome of TBAV and TTAV aortas, with the same genetic background but different valve morphology, we identified 110 differentially expressed proteins, many of which belong to pathways related to cystic medial degeneration (Supplemental Table 10), with several pathways associated with shear stress due to altered hemodynamics in TAD patients^2,59,60^. Of these 110 differentially expressed proteins, 55.2% were concordant with the comparison between TBAV and HTAV, which includes the genetic plus the hemodynamic components. Indeed, both the heatmap and the enrichment analyses of TBAV *vs.* TTAV were clearly more similar to TBAV *vs.* HTAV than to TTAV *vs.* HTAV (compare in Figs. 1C and 2). All these findings indicate that hemodynamic disturbances caused by the valve morphology, rather than the genetic background, are the main inducers of most molecular alterations in the aorta of BAV individuals. It can then be proposed that genetic factors predispose to BAV aortopathy, whereas hemodynamics determine the progression of the disease. This hypothesis agrees with the observed higher prevalence of aortic dilatation in children with valve disfunction^61,62^.

Currently, the severity of BAV aortopathy that establishes the necessity of surgery is determined by the diameter of the aorta, although there are no reference values that serves as a concrete cutoff. Several studies have attempted to find diagnostic or prognostic biomarkers^63^, but no feasible biomarkers have been identified until now^16^. The IPA biomarker filter used in this study identified 23 potential biomarkers (overexpressed in TBAV specimens) detectable in sentinel tissues (i.e., blood, peripheric blood mononuclear cells, urine, and other bodily fluids). Through literature research, we found four additional differentially expressed proteins in TBAV animals, detectable in blood samples, which were previously identified in TAD patients (Supplemental Table 11). Therefore, we present 27 new potential biomarkers with a high potential utility (Table 1). Considering that the hamster T-strain serves as a model of BAV aortopathy predisposition, exhibiting the histopathologic substrate and TAD but not aneurysm, the proposed molecular biomarkers may have predictive value as diagnostic markers.

**Table 1.** Proposed new potential biomarkers. Proteins with differential expression in TBAV animals, detectable in sentinel tissues.

| Symbol | Entrez Protein Name | p-value | Fold Change | Type |
| --- | --- | --- | --- | --- |
| **TIMP2** | Metalloproteinase inhibitor 2 | - | H<br>Exclusive | peptidase |
| **VCL** | Vinculin isoform X1 | - | H<br>Exclusive | other |
| **HIBADH** | 3-hydroxyisobutyrate dehydrogenase | 0.0289 | -5.02 | enzyme |
| **ROCK1** | Rho associated coiled-coil containing protein kinase 1 | 0.0401 | -3 | kinase |
| **RASL12** | RAS like family 12 | 0.0018 | -2.577 | enzyme |
| **RPL17** | ribosomal protein L17 | 0.0175 | -2.351 | other |
| **LGMN** | Legumain | 0.0261 | -2.184 | protease |
| **PRKAA1** | protein kinase AMP-activated catalytic subunit alpha 1 | 0.00524 | -1.979 | kinase |
| **AK1** | adenylate kinase 1 | 0.0203 | -1.974 | kinase |
| **AEBP1** | AE binding protein 1 | 0.0335 | -1.939 | peptidase |
| **VTN** | vitronectin | 0.0415 | -1.785 | other |
| **CSRP2** | cysteine and glycine rich protein 2 | 0.00727 | -1.684 | other |
| **PSMD8** | proteasome 26S subunit, non-ATPase 8 | 0.0138 | -1.651 | other |
| **HSPA5** | heat shock protein family A (Hsp70) member 5 | 0.0428 | -1.378 | enzyme |
| **FBLN5** | fibulin-5 isoform | 0.0117 | -1.054 | other |
| **SEC22B** | SEC22 homolog B, vesicle trafficking protein | 0.0495 | -0.985 | other |
| **RAB21** | RAB21, member RAS oncogene family | 0.0177 | 1.002 | enzyme |
| **TBCC** | tubulin folding cofactor C | 0.0278 | 1.13 | enzyme |
| **GNPDA1** | glucosamine-6-phosphate deaminase 1 | 0.0417 | 1.539 | enzyme |
| **GC** | GC vitamin D binding protein | 0.0462 | 1.558 | transporter |
| **ACSS3** | acyl-CoA synthetase short chain family member 3 | 0.0461 | 1.626 | enzyme |
| **ATP6V1A** | ATPase H<sup>+</sup> transporting V1 subunit A | 0.00152 | 1.655 | enzyme |
| **TMED2** | transmembrane p24 trafficking protein 2 | 0.015 | 1.827 | other |
| **SOD1** | superoxide dismutase 1 | 0.0064 | 1.852 | enzyme |
| **SLIT3** | slit guidance ligand 3 | 0.016 | 2.553 | other |
| **GOLGA3** | golgin A3 | 0.00223 | 2.927 | other |
| **RPL18A** | ribosomal protein L18a | 0.0208 | 3.843 | other |

### Study limitations

This is a hypothesis-generating study based on the comparison of an animal model with the patient population. As such, one obvious limitation is the possible influence of interspecific differences. However, these differences do not impact the principal conclusion of the study, i.e., the existence of a set of interacting altered pathways that characterizes TAD, because this was precisely deduced from similarities between the two species.

One intrinsic constrain of the model is that animals show the histopathological and molecular substrate of the disease, but do not develop aortopathy, making it a predisposition model. The T-strain may represent a patient population with a high predisposition to BAV and TAD development, but less prone to aneurysm. Alternatively, absence of aneurysm in hamsters may result from species-specific insufficient physiopathological progression of the disease, including their short average life expectancy, or possible biomechanical differences with humans due to the very small size of the hamster aorta.

## Summary and Conclusions

Based on our results from genetically and phenotypically homogeneous animal model populations, as well as multiple studies involving different and heterogeneous patient populations, we propose the existence of a common assemblage of interconnected altered molecular pathways shared by individuals with BAV-associated TAD. These pathways, NF-κB, WNT/CTNNB1, mTOR, MAPK1/2 and eNOS are directly regulated by the central PI3K/AKT pathway, the overactivation of which depends on alterations of TGF-β, ANGII and EGF effectors. WNT/CTNNB1 signaling may be a major genetic factor contributing to BAV disease. Alterations in this pathway may induce phenotypic shift specifically in CNC-derived SMCs, favored by oxidative stress and inflammation. We propose that while genetic factors cause the structural abnormalities of the aorta in TAD individuals, hemodynamics are the main triggers of proteome alterations. Therefore, genetic factors predispose to pathology whereas hemodynamics determine progression. Finally, our study reveals 27 new potential biomarkers with predictive and/or diagnostic value.

## Acknowledgments

We acknowledge Adrian Ruíz Villalva for technical and scientific support, Luis Vida for technical assistance, Francisco David Navas and Jessica Román Pérez for assistance in confocal microscopy, and Remedios Crespillo Molina for assistance in molecular techniques.

## Abbreviations list

α-Actin: Alpha smooth muscle
AKT: Protein kinase B
ANGII: Angiotensin-2
ATR-1: Angiotensin II receptor type 1
BAV: Bicuspid aortic valve
CNC: Cardiac Neural Crest
CSDE: Cold Shock Domain-containing E
CTNNB1: Beta-catenin1
DAF-2DA: 4,5-diaminofluorescein diacetate
EGF: Epidermal growth factor
EGFR: Epidermal growth factor receptor
ERBB: Receptor tyrosine-protein kinase
EMT: Epithelia-to-mesenchymal transition
eNOS: Endothelial nitric oxide synthase
ERK: Extracellular signal-regulated kinases
Fbln: Fibrillin
GSk3b: Glycogen synthase kinase 3 beta
HTAV: TAV animals from the H-strain
IF: Immunofluorescence
IKK: IkappaB kinase
IPA: Ingenuity pathway analysis software
MAPK: Mitogen-activated protein kinases
mTOR: Mammalian Target of Rapamycin
NF-κB: Nuclear factor κB
NO: Nitric oxide
NOTCH: Neurogenic locus notch homolog protein
pAKT: Activated AKT
PI3K: Phosphoinositide 3-kinase
PPI: Protein-protein interaction
pSMAD2: Activated SMAD2
RAF: Rapidly Accelerated Fibrosarcoma
RAS: Rat sarcoma virus
RHEB: Ras homolog enriched in brain
SMAD2: Mothers Against Decantaplegic homolog 2
SMC: Smooth muscle cell
TAD: Thoracic aortic dilatation
TAV: Tricuspid aortic valve
TBAV: BAV animals from the T-strain
TGF-β: Transforming growth factor-beta
TIMP: Metallopeptidase inhibitor
TTAV: TAV animals from the T-strain
WB: Western blot
WNT: Wingless-related integration site

